# Hematopoietic stem cell requirement for macrophage regeneration is tissue-specific

**DOI:** 10.1101/2021.04.08.439077

**Authors:** Devon J. Eddins, Astrid Kosters, Jeffrey Waters, Jasmine Sosa, Megan Phillips, Koshika Yadava, Leonore A. Herzenberg, Hedwich F. Kuipers, Eliver Eid Bou Ghosn

**Author notes:** Joint first authors (Devon J. Eddins, Astrid Kosters). ***Corresponding author contact info***, Eliver E.B. Ghosn: Lowance Center for Human Immunology, Health Sciences Research Building, 1760 Haygood Dr. NE, E240, Atlanta, GA 30322, USA.

## Abstract

Tissue-resident macrophages (TRMΦ) are important immune sentinels responsible for maintaining tissue and immune homeostasis within their specific niche. Recently, the origins of TRMΦ have undergone intense scrutiny where now most TRMΦ are thought to originate early during embryonic development independent of hematopoietic stem cells (HSCs). We previously characterized two distinct subsets of mouse peritoneal cavity macrophages (Large and Small Peritoneal Macrophages; LPM and SPM, respectively) whose origins and relationship to both fetal and adult long-term (LT)-HSCs have not been fully investigated. Here we employ highly purified LT-HSC transplantation and in vivo lineage tracing to show a dual ontogeny for LPM and SPM, where the initial wave of peritoneal macrophages is seeded from yolk sac-derived precursors, which later require LT-HSCs for regeneration. In contrast, transplanted fetal and adult LT-HSCs are not able to regenerate brain-resident microglia. Thus, we demonstrate that LT-HSCs retain the potential to develop into TRMΦ, but their requirement is tissue-specific.

## Introduction

Virtually all known organs in vertebrates contain tissue-resident macrophages (TRMΦ) that serve important roles in maintaining tissue and immune homeostasis therein (Davies et al., 2013a; Li and Barres, 2018; Wynn et al., 2013). It was long assumed that all macrophages (MΦ) develop from monocytes generated by hematopoietic stem cells (HSCs) in the bone marrow (BM) (Osawa et al., 1996; Smith et al., 1991; Till and McCulloch, 1980; van Furth and Cohn, 1968). However, in recent years, an overwhelming body of evidence has overtly challenged the notion that TRMΦ are solely derived from HSCs (Ginhoux et al., 2010; Gomez Perdiguero et al., 2015a; Yona et al., 2013). Collectively, these studies have established that most TRMΦ populations develop during embryogenesis from yolk sac progenitors that emerge prior to, and independent of, long-term (LT)-HSCs. Tissue-resident macrophages, including brain microglia (Ginhoux et al., 2010) and skin Langerhans cells (Gomez Perdiguero et al., 2015a; Hoeffel et al., 2015), emerge at around embryonic day 8 (E8) in a region of the yolk sac known as the blood island, before the development of the first definitive LT-HSC (which starts at E10.5) (Ghosn et al., 2019). Before birth, these yolk sac-derived MΦ migrate and take long-term residence in the various tissues (i.e., brain and skin) of the developing embryo. Fate-mapping (Buttgereit et al., 2016; Ginhoux et al., 2010; Gomez Perdiguero et al., 2015a; Yona et al., 2013) and parabiosis (Ajami et al., 2007; Hashimoto et al., 2013; Huang et al., 2018) experiments show that TRMΦ are maintained throughout adulthood by *in situ* self-renewal, with minimal contribution from LT-HSC-derived circulating monocytes.

We previously identified and characterized two functionally distinct subsets of TRMΦ in the mouse peritoneal cavity (PerC), namely Large and Small Peritoneal Macrophages (LPM and SPM, respectively) (Ghosn et al., 2010). In subsequent years, various studies demonstrating that most TRMΦ (i.e., microglia, Kupffer cells, Langerhans cells, etc.) develop from HSC-independent, yolk sac-derived fetal progenitors (Gomez Perdiguero et al., 2015a), have led some researchers to speculate that LPM and SPM are also likely to develop independently of LT-HSCs (Cassado et al., 2015). Although peritoneal macrophages are one of the most studied TRMΦ populations, with much previously done to identify the functional and developmental differences between LPM and SPM (Bain et al., 2016; Broche and Tellado, 2001; Cain et al., 2013; Ghosn et al., 2010), their origins and relationship to both fetal and adult LT-HSCs have not been fully investigated and remains controversial. Despite the mounting evidence supporting the notion that TRMΦ originate from HSC-independent yolk sac-derived progenitors (Gomez Perdiguero et al., 2015a; Gomez Perdiguero et al., 2015b) and/or fetal liver monocytes (Hoeffel et al., 2012; Hoeffel et al., 2015), opposing studies have suggested that all TRMΦ, with the exception of microglia and a fraction of Langerhans cells, are instead derived from fetal LT-HSCs (Sheng et al., 2015a, Sheng et al., 2015b). Although these two hypotheses are not necessarily mutually exclusively, they have not been tested simultaneously. Most importantly, the potential of highly purified and transplanted *bona fide* LT-HSCs, from both fetal and adult sources, to fully regenerate TRMΦ *in vivo* (including LPM, SPM, and microglia) has not been tested.

To resolve these seemingly contradictory findings and determine whether certain TRMΦ show single or dual ontogeny, we directly tested the potential of highly purified fetal and adult LT-HSCs to regenerate TRMΦ in the peritoneum and brain of lethally-irradiated recipient mice. We show that both fetal and adult LT-HSCs fully regenerate tissue-resident LPM and SPM populations, but completely fail to regenerate tissue-resident microglia. On the other hand, using *Runx1* lineage-tracing, we show that, similar to brain microglia, E8 progenitors can also give rise to both LPM and SPM independently of LT-HSCs. In conclusion, our studies show a dual ontogeny for tissue-resident LPM and SPM (i.e., HSC-independent and HSC-dependent), and confirm the HSC-independent origin for brain microglia. Importantly, we directly demonstrate that brain microglia, unlike LPM and SPM, cannot be regenerated by transplantation of purified *bona fide* fetal LT-HSCs, even after the host received lethal, full-body irradiation. Collectively, these findings add a new layer to the complex developmental landscape of the myeloid lineage and challenge the current notion that LPM and SPM have divergent and distinct origins.

## Materials and Methods

### Mice and Tissue Preparation

C57BL/6 and Gt(ROSA)26Sor^tm4(ACTB-tdTomato,-EGFP)Luo^/J (ROSA^mT/mG^, Stock No: 007576) mice (8-10 wks) were purchased from Jackson Laboratory (Bay Harbor, ME, USA). Transgenic mice expressing enhanced green (pCx-eGFP) (Wright et al., 2001) or red (TM7-RFP) (Ueno and Weissman, 2006) fluorescent protein were kindly provided by the Weissman laboratory (Stanford). *Runx1*^MerCreMer^ mutant mice were generated by Dr. Igor Samokhvalov and colleagues (Samokhvalov et al., 2007) and provided by Riken Center for Life Science Technologies (accession number CDB0524K; http://www.clst.riken.jp/arg/mutant%20mice%20list.html; Wako, Saitama Prefecture, Japan). Mice were housed and bred at Emory and Stanford animal facilities. *Runx1*^MerCreMer^ mice were crossed with ROSA^mT/mG^ mice to generate tamoxifen-inducible fate mapping model (*Runx1*^cre/eGFP^). Timed pregnancies were confirmed via post-coital plug and embryonic ages were confirmed via microscopy. Peripheral blood from progeny mice was screened for multi-lineage eGFP^+^ cells to ensure LT-HSCs were not labeled prior to being used in experiments. All procedures were approved by both Emory and Stanford Institutional Animal Care and Use Committees (IACUC) in compliance with the recommendations in the Guide for the Care and Use of Laboratory Animals of the National Institutes of Health and follow administrative panel on laboratory animal care (APLAC) guidelines.

Blood (~200 μL) was drawn via tail vein into EDTA-containing tubes (BD Diagnostics). Peritoneal cavity (PerC) cells were harvested by PerC lavage with 7 mL of custom RPMI-1640 media deficient in biotin, L-glutamine, phenol red, riboflavin, and sodium bicarbonate, with 3% newborn calf serum and benzonase (defRPMI). Bone marrow (BM) from femurs and tibias were flushed with defRPMI using 28G needle and 30 mL syringe (BD Medical). Cells were passed through a 70 μm nylon filter (Corning) and erythrocytes were lysed using ACK lysis buffer. Whole brains from mice were mechanically dissociated using a Dounce homogenizer then gently passed through a 70 μm filter and centrifuged at 300 g, 4°C, 10 min. Cells were resuspended in a 28% isotonic Percoll cushion (GE Healthcare) and centrifuged (900 g, 4°C, 30 min.) to remove myelin. Fetal livers were harvested from ≥E15 timed pregnant mice, digested at 37°C for 30 min. with 0.25% collagenase I (Stem Cell Technologies), then again using enzyme-free cell dissociation buffer (Gibco). Liver cells were passed through 70 μm filter to obtain single cell suspensions.

### Tamoxifen treatment

Homozygous male ROSA^mT/mG^ mice were mated with heterozygous female *Runx1*^MerCreMer^ mice overnight (at 18:00). Female mice were examined for post-coital plug the following morning (at 08:00), and those with a plug were considered pregnant and timed E0.5. Pregnant mice received a single dose of 0.1 mg/g body weight (Z)-4-Hydroxytamoxifen supplemented with 0.05 mg/g body weight progesterone resuspended in corn oil (all from Sigma-Aldrich) via intraperitoneal (i.p.) injection at E8. Progesterone supplementation counteracts estrogen receptor antagonism by tamoxifen to circumvent fetal abortions.

### 18-parameter High-Dimensional (Hi-D) Flow Cytometry

Cells were resuspended at ≤1 x 10^7^ cells/mL in defRPMI and stained on ice for 30 min. (or 60 min. when staining for CD34) with the following fluorochrome-conjugated mAbs (see Table S1 for clones and sources). Briefly, for recipient PerC: anti-CD5, CD19, F4/80, NK1.1, IgM, IgK, VH11, CD23, I-A/I-E, CD11b, Gr-1, CD45, and B220; Brain: anti-CD11b, F4/80, CX3CR1, CD5, CD19, CD11c, CD45, CD23, Gr-1, Ly-6C, CD43, I-A/I-E, and CD16/32; and blood: anti-TER-119, CD5, CD45, Ly-6C, Gr-1, CD11b, NK1.1, and CD19. Cells were stained on ice for 15 min with Qdot605- or BV711-conjugated streptavidin to reveal biotin-coupled antibodies (see Table S1). Stained cells were re-suspended in 10 μg/mL propidium iodide (PI) to exclude dead cells. When cells were fixed, amine-reactive dyes were used to exclude dead cells. Both GFP and RFP were detected concomitantly with the reagents described above for a total of 18-parameter Hi-D FACS. Cells were analyzed on Emory Pediatric/Winship Flow Cytometry Core or Stanford Shared FACS Facility instruments (BD LSRII). Data were collected for 0.2–3 x 10^6^ cells and analyzed with FlowJo (FlowJo LLC). To distinguish auto-fluorescent cells from cells expressing low levels of a particular surface marker, we established upper thresholds for auto-fluorescence by staining samples with fluorescence-minus-one (FMO) control stain sets in which a reagent for a channel of interest is omitted.

### Sorting and transfer

Fetal liver from ~E15 TM7-RFP (RFP^+^) mice were processed as described above and stained with the following mAbs in a 15-color, 17-parameter staining combination: anti-SCA-1, CD38, CD150, CD34, CD48, CD117, CD41, CD45, CD127, CD135, CD19, and lineage markers (Lin) anti-CD3ε, B220, NK1.1, Gr-1, CD11b, TER-119 (Table S1). Cells were stained on ice for 15 min with BV711-conjugated streptavidin to reveal biotin-coupled mAbs and re-suspended in PI, to exclude dead cells. RFP^+^ or GFP^+^ LT-HSCs were identified as Lin^-^, CD117^hi^, SCA-1^hi^, CD150^+^, CD48^-^, CD41^-^, CD34^low^, CD45^+^, CD38^+^, CD127^-^, and CD135^-^ (Fig. S4). Sorted LT-HSCs were re-suspended in serum free defRPMI and about 100 cells were transferred intravenously (i.v.) into lethally irradiated (two doses of 4.25 Gy delivered 4 h apart) C57BL/6 mice along with ~2 x 10^5^ BM rescue cells from 8 wks old congenic pCx-eGFP (eGFP^+^) mice. After 30+ wks, recipient PerC, Brain, and blood cells were harvested and LT-HSC-derived TRMΦ (RFP^+^) analyzed as described above. To determine TRMΦ reconstitution potential for adult BM LT-HSCs, BM from RFP^+^ mice were processed and stained as described for fetal liver. LT-HSCs were sorted on Emory Pediatric/Winship Flow Cytometry Core or Stanford Shared FACS Facility BD FACSAria II instruments.

### Chimerism

Lethally irradiated recipient C57BL/6 (10-20 wks) mice were injected i.v. with either ~100 sorted fetal liver (~E15 embryos) or adult BM (>20 wks) LT-HSCs from donor RFP^+^ mice, along with ~2 x 10^5^ adult BM rescue cells from congenic eGFP^+^ mice. Co-transfer of the rescue cells is necessary to prevent the lethally irradiated mice from succumbing from anemia during the first weeks post-irradiation. It readily provides red and white blood cells until the transferred LT-HSCs reconstitute all major blood cells. Blood was collected weekly from recipient mice to determine the level of chimerism, which we defined as the percentage of cells derived from the donor fetal liver or adult BM LT-HSCs (RFP^+^) found among total blood cells recovered from recipient mice. Here, we examined the recipient mice that showed full, long-term stable chimerism (i.e., development of all major hematopoietic lineages, including erythrocytes, myeloid cells, granulocytes, T cells, and B cells, see Fig. S5). All the reconstituted hematopoietic lineages from donor cells were still readily detectable in recipient blood when the mice were sacrificed, and tissues harvested at ≥33 wks post transplantation (Fig. S5). A total of 40 recipient mice received LT-HSC transplantation (13 adult BM and 27 fetal liver) in seven independent transplantation experiments. Out of 40 mice, 26 became chimeric (11 adult BM and 15 fetal liver) and we chose the cohorts with the highest blood chimerism (4 adult BM and 11 fetal liver, see Fig. S5). The data shown in the various figures represent the fifteen fully chimeric mice that received purified LT-HSCs, and the FACS plots shown in each figure represent the analysis from the same recipient mouse.

### Fluidigm single-cell multiplexed qPCR

Single-cell multiplexed qPCR experiments were performed using Fluidigm’s (San Francisco, CA, USA) 96.96 qPCR DynamicArray microfluidic chips as previously described (Lawson et al., 2015). PerC cells were stained with anti-F4/80, CD5, CD19, NK1.1, Gr-1, CD11b, and I-A/I-E, then single cells were FACS-sorted into individual wells of 96-well PCR plates, using the FACSAriaII. Experiments were performed following Fluidigm’s Advanced Development Protocol 41. The 96-well plates w preloaded with 9 μL of RT-STA solution: 5 μL of CellsDirect PCR mix (Invitrogen), 0.2 μL of SuperScript-III RT/Platinum Taq mix (Invitrogen), 1.0 μL of a mixture of all pooled primer assays (500 nM), and 2.8 μL of TE buffer (Promega). After sorting, PCR plates were either frozen (−80°C) or immediately run for reverse transcription (50°C for 15 min, 95°C for 2 min) and target-specific amplification (20 cycles; each cycle: 95°C for 15 s, 58°C for 4 min). Biological replicates were performed in lieu of technical replicates per the manufacturer’s recommendation, to yield more power and better sampling of the target populations. 3.6 μL of exonuclease reaction solution (2.52 μL sterile nuclease-free water, 0.36 μL Exo reaction buffer, and 0.72 μL Exo-I, New England BioLabs) was then added to remove unincorporated primers (37°C for 30 min, 80°C for 15 min), then each well was diluted 1:3 with TE buffer (Promega). A 2.7 μL aliquot from each sample was mixed with 2.5 μL of SsoFast EvaGreen Supermix with Low Rox (Bio-Rad) and 0.25 μL of Fluidigm’s DNA Binding Dye Sample Loading Reagent in a separate plate and centrifuged to mix solutions. Individual primer assay mixes were generated in each well of a separate plate by loading 2.5 μL of Assay Loading Reagent (Fluidigm), 2.25 μL DNA Suspension Buffer (TEKnova), and 0.25 μL of 100 μM primer pair mix. Chips were primed by injecting control line fluid (Fluidigm) into each accumulator on the integrated fluidics circuit (IFC) and running the ‘Prime’ program prior to loading primer assays and samples. 5 μL of each sample and primer mix were loaded into each well of the chips. Samples and assays were then mixed in the chip by running the ‘Load Mix’ program in the IFC Controller HX. Chips were loaded into the BioMark real-time PCR reader (Fluidigm) and run following the manufacturer’s protocol. A list of primer assays used in this study is provided in Table S2.

### In vivo phagocytosis

Chimeric mice received 500 μL i.p. injections of 1 mg/mL pHrodo-labeled *Escherichia coli* particles (Thermo Fisher Scientific) resuspended in defRPMI-1640. Peritoneal macrophages were isolated via PerC lavage 2 h after i.p. injection with pHrodo-labeled *E. coli*, then analyzed by Hi-D flow cytometry (as described above) and fluorescent microscopy.

### Fluorescent microscopy

LPM/SPM isolated from mice 2 h after receiving pHrodo-labeled *E. coli* injections were stained with anti-F4/80 (AF488), CD19, CD11c, I-A/I-E, and CD11b, then FACS sorted directly into an 8-well chamber slide in a live cell imaging solution. Cells were mounted with a ProLong® diamond antifade mountant with DAPI (Thermo Fisher Scientific) and imaged using a Leica SP5 multiphoton/confocal Laser Scanning microscope at Stanford’s Cell Sciences Imaging Facility.

### Statistical analyses

All graphing and statistical analyses were performed using GraphPad Prism v9. Unpaired t-tests were used to determine statistical differences (p<0.05) where indicated. Data were analyzed for distribution (normal (Gaussian) vs. lognormal) independently using the Shapiro-Wilk test for normality in both the untransformed and Log10 transformed data. When data passed both distribution tests, the likelihood of distribution (normal vs. lognormal) was computed and QQ-plots generated for both untransformed and Log10 transformed data. When Log10 transformed data had a higher likelihood of a normal distribution (passing normal distribution test) and/or failed lognormal distribution test, parametric analyses were performed. If the data had unequal variance (as determined by a F test on both the untransformed and Log10 transformed data), Welch’s T test was performed. All instances where lognormal distribution was more likely non-parametric (Mann-Whitney) tests were performed.

## Results

### Tissue-resident macrophages develop and take residence during early fetal development

To investigate the moment in development that myeloid cells populate PerC and brain tissues, we determined the presence of immune cells in these compartments throughout embryonic and postnatal development in mice. This analysis demonstrated that both PerC and brain contain immune cells as early as E10, preceding development of the first LT-HSC, which has been shown to start at E10.5 (Kieusseian et al., 2012; Kumaravelu et al., 2002; Muller et al., 1994), and that these first immune cells are almost exclusively CD11b^+^ myeloid cells (Fig. 1). Throughout prenatal development, the majority of the immune cells in the PerC are myeloid (>85%). However, during peri- and postnatal development, other immune cells (CD45^+^, CD11b^lo^, F4/80^−^) populate this tissue as well, resulting in ~20% of peritoneal cells being myeloid in the adult mouse (Fig. 1). Finally, we observed that in both newborn and adult animals, the majority of peritoneal MΦ are LPM (95% and 90%, respectively). Conversely, microglia remain the predominant leukocyte population in the brain throughout life. Together, these data indicate that the first wave of TRMΦ emerge and colonize tissues prior to the development of LT-HSCs. In addition, they demonstrate that the MΦ compartment in adult animals differs between peritoneal and brain tissue.

**Figure 1.**
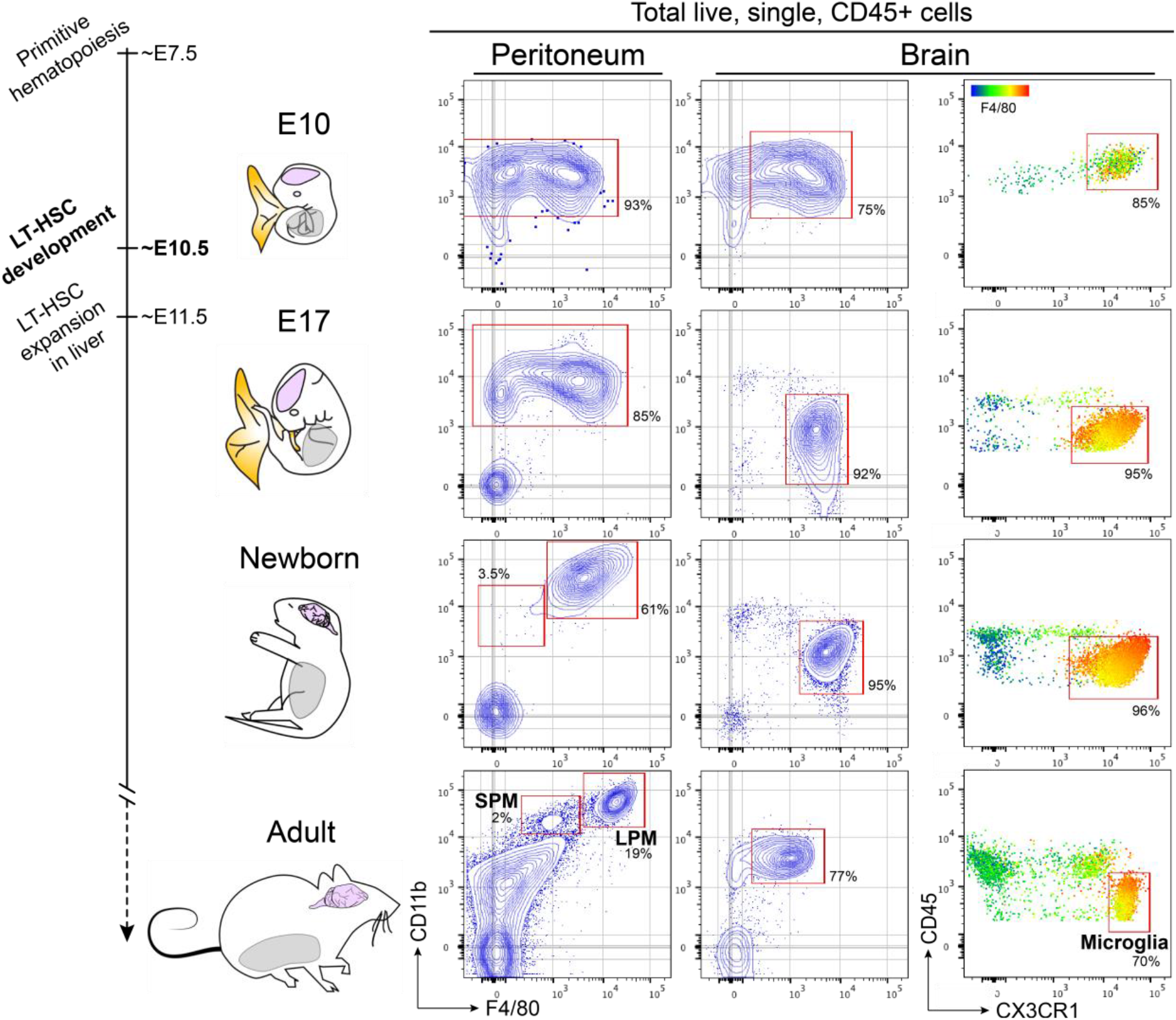
Tissue-resident peritoneal and brain macrophages develop and take residence during early fetal development. The composition of total peritoneal and brain tissues from mice at various developmental ages, analyzed by flow cytometry. The percentages of myeloid cells, identified as CD11b^+^ (peritoneum and brain) and/or CX3CR1^+^ (brain), are shown in these tissues at embryonic day 10 (E10), embryonic day 17 (E17), 2 days after birth (newborn) and >8 weeks (adult). SPM and LPM were additionally distinguished by F4/80 expression in peritoneal tissue of newborn and adult animals. Data shown are representative of >10 mice in 3 independent experiments. All cells shown were pre-gated to include only live, single cells, expressing CD45.

### Tissue-resident macrophages derive from an early myeloid progenitor separate from the fetal LT-HSC

The observation that myeloid cells are already present in peritoneal and brain tissue before the first LT-HSC develops infers that they are derived from a separate, LT-HSC-independent source. Therefore, we next determined the fetal origin of tissue resident myeloid cells. For this, we employed a lineage tracing method, using inducible *Runx1*^cre/eGFP^ (*Runx1*^MerCreMer^) mice to label yolk sac progenitors (Samokhvalov et al., 2007) and track their progeny into adult life. We labeled fetal Runx1^+^ immune progenitor cells at E8, at which time they have been shown to be located in the yolk sac (Samokhvalov et al., 2007) and which is well before the first LT-HSC develop (Fig. 2A). A fraction of both peritoneal (~10%) and brain (~6%) macrophages in animals are derived from yolk sac progenitors that were labeled at E8 (Fig. 2D) and were maintained until adulthood (Fig. S1). Furthermore, the distribution of labeled, E8 progenitor-derived MΦ in the peritoneum is very similar to that of unlabeled MΦ (i.e., the LPM/SPM ratio within GFP^+^ and GFP^-^ compartments), suggesting that the composition of these populations is stable throughout adult life, where the majority of peritoneal MΦ are LPM (see Fig. 2D).

**Figure 2.**
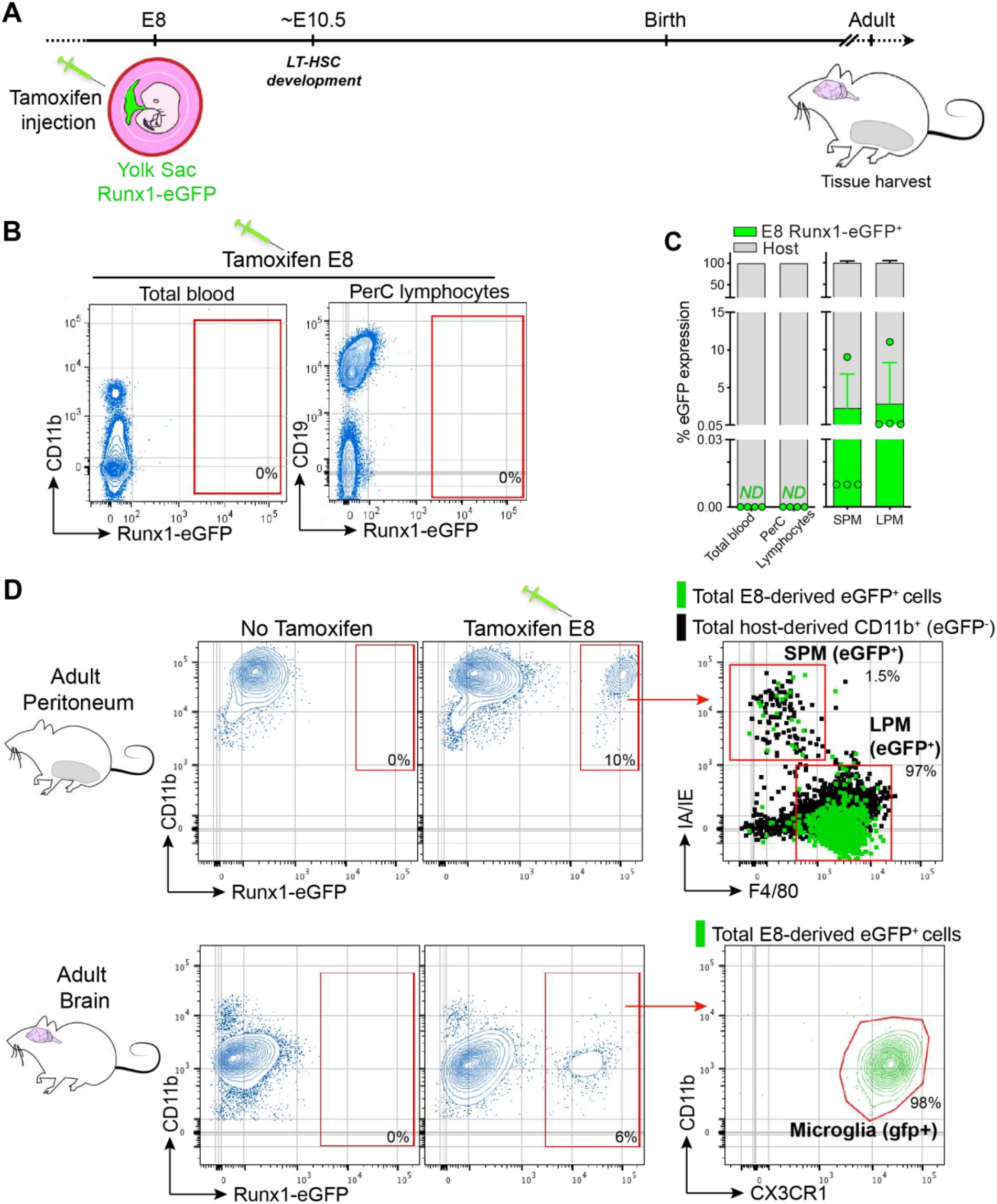
Tissue-resident SPM, LPM, and microglia emerge before the development of fetal LT-HSC. **A.** Schematic overview of the lineage-tracing assays to track the progeny of fetal (E8) progenitor cells. Tamoxifen was injected into pregnant *Runx1*^MerCreMer^ x ROSA26^mT/mG^ mice, then peritoneum and brain were analyzed for eGFP expression in adult offspring (*Runx1*^Cre/eGFP^). **B.** Analysis of eGFP expression in circulating cells, as well as in lymphocytes in the peritoneum, of adult mice, showing that E8 tamoxifen injection did not label hematopoietic stem cells and other lymphoid lineages. **C.** Quantification of eGFP signal in peripheral blood and PerC lymphocytes (B) and SPM/LPM (D) of *Runx1*^Cre/eGFP^ mice. **D.** Flow cytometry analysis of tissue-resident macrophages (TRMΦ) (CD11b^+^) in the peritoneum and brain of adult mice, showing the percentage of TRMΦ that are derived from E8-labeled (GFP^+^) progenitors. Data shown are representative of 4 mice from 3 experiments. See Figs. S2 and S3 for representative gating strategies of PerC and brain. ND = not detected.

To confirm that the E8-labeled progenitor-derived TRMΦ were not derived from fetal LT-HSC, we assessed whether any circulating LT-HSC-derived hematopoietic cells were labeled in adult animals. Assessing eGFP expression in the blood and in lymphocyte populations in the PerC confirmed the absence of multi-lineage labeling (Fig. 2B), indicating that the cells labeled at E8 were not LT-HSCs, but indeed a separate myeloid progenitor, giving rise exclusively to TRMΦ. These data demonstrate that, similar to microglia, tissue-resident LPM and SPM are initially derived from a myeloid progenitor that emerges early during embryogenesis in the yolk sac and is separate from the fetal LT-HSC.

### Fetal and adult LT-HSC transplantation reveals a dual ontogeny for tissue-resident peritoneal macrophages, but not brain microglia

After determining that myeloid progenitors present in the yolk sac at E8 give rise to TRMΦ, we next asked whether LT-HSCs, which emerge at E10.5, retain the potential to generate these TRMΦ. To this end, we highly purified LT-HSCs from fetal liver (at E15) and adult bone marrow (at 24 wks of age) and transplanted ~100 purified cells into lethally irradiated mice (Fig. S4). We then analyzed the progeny of these transplanted cells in tissues of fully chimeric mice at least 33 wks after transplantation (Fig. 3A,B,S5). This analysis revealed that purified LT-HSCs from either source (fetal liver or adult BM) represent true HSCs, capable of generating all components of the immune system (Fig. 3A,S5). LT-HSCs from fetal liver exhibit a reconstitution advantage over host and rescue BM cells, which has been previously reported (Ghosn et al., 2016; van de Laar et al., 2016; Fig. 3A,S5). Moreover, both fetal and adult LT-HSCs are capable of readily generating both LPM and SPM in the peritoneum (Fig. 3C,E). However, neither fetal nor adult LT-HSCs regenerate microglia in the adult brain, even after lethal, full-body irradiation of the recipient animals (Fig. 3D,F). In contrast, we confirmed that monocytes and other blood-derived immune cells that were present in the brain preparations are derived from LT-HSCs (Fig. 3D,F), suggesting that microglia residing in the brain parenchyma are maintained independent of fetal and adult LT-HSCs or circulating monocytes. Together, these data indicate that TRMΦ in the adult peritoneum, but not in the brain, require LT-HSCs for regeneration.

**Figure 3.**
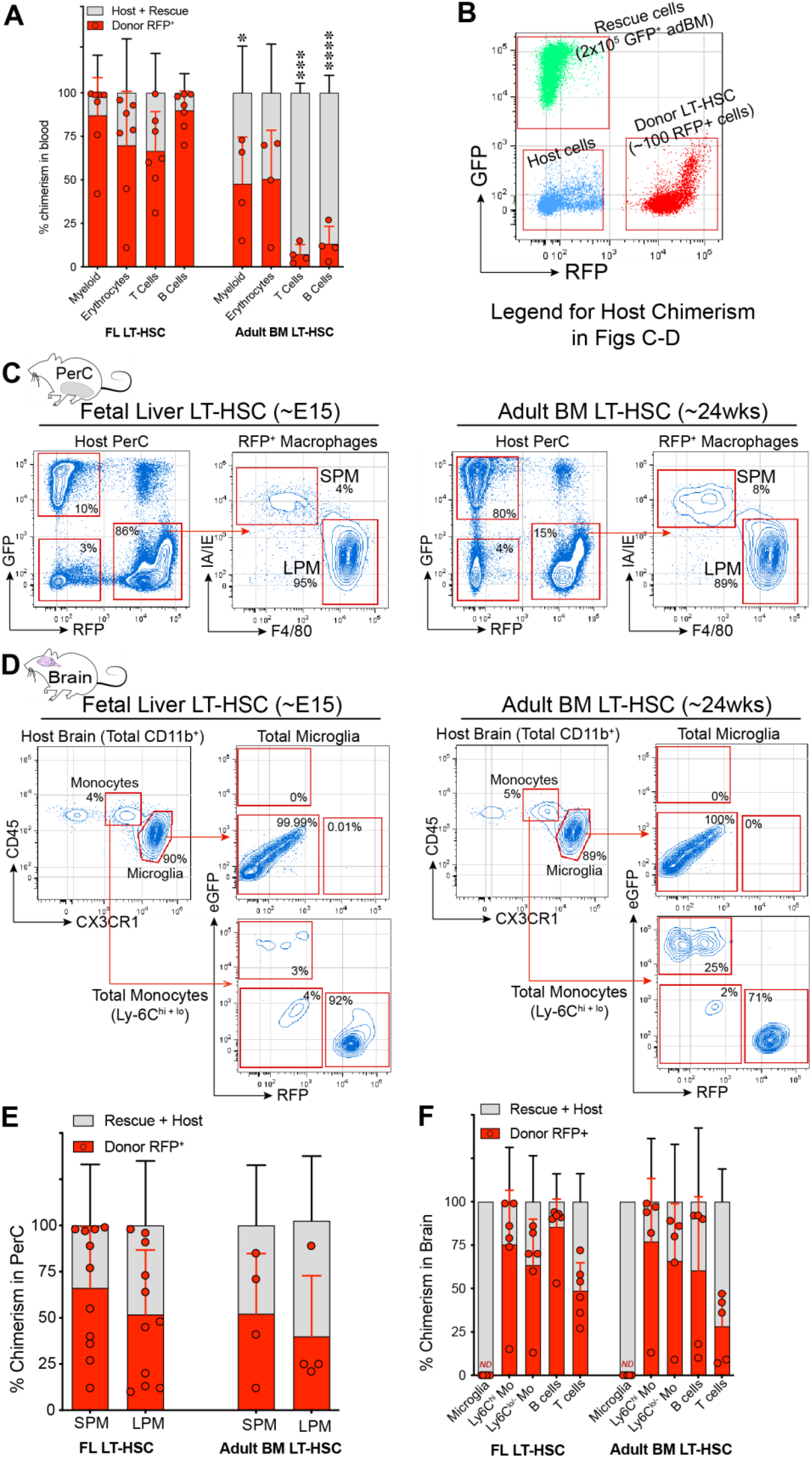
Transplanted LT-HSC from fetal and adult origin fully regenerate tissue-resident peritoneal macrophages, but not brain microglia. **A.** Blood chimerism rates of transplanted highly purified LT-HSC that were isolated from fetal liver (FL) or adult bone marrow (BM). Shown are the fraction of cells that were derived from transplanted (RFP^+^) LT-HSC, in different immune compartments in the blood of adult animals. **B.** Overview of the analysis/gating strategy to determine the progeny of transplanted purified LT-HSC. **C–D.** Analysis of RFP expression in **C**) SPM and LPM in the peritoneal cavity and **D**) microglia and monocytes in the brain of adult animals that were transplanted with purified LT-HSC that were isolated from fetal liver or adult bone marrow. **E.** Chimerism rates of transplanted (RFP^+^) fetal and adult LT-HSC-derived cells among SPM and LPM in the peritoneal cavity. **F.** Chimerism rates of transplanted (RFP^+^) fetal and adult LT-HSC-derived cells among microglia, and other immune cell subsets in the brain of recipient mice. Data shown are mean + SD (A, E and F) and representative (C and D) of the 15 fully chimeric animals (4 adBM and 11 fetal). Comparisons between LT-HSC source * p = 0.0255 (unpaired t-test), *** p = 0.0003 (Welch’s t-test), **** p = <0.0001 (unpaired t-test). ND = not detected, red points = RFP chimerism of individual replicates.

### Tissue-resident peritoneal macrophages derived from LT-HSC transplants are functionally comparable to their naturally occurring counterpart

Because the TRMΦ in the peritoneum can regenerate from LT-HSCs, in addition to being derived from early HSC-independent myeloid progenitors, we assessed whether peritoneal macrophages derived from LT-HSC transplantation are functionally different from their naturally developed (native) counterparts. To test this, we performed a targeted single-cell transcriptomics assay (multiplex qPCR), assessing the expression of a curated set of 77 transcription factors, on single-captured MΦ (Fig. 4A) isolated from the PerC of adult mice that were fully chimeric for transplanted fetal RFP^+^ LT-HSCs. Analyzing the expression profile of this set of transcription factors allows for sensitive classification of cell types and hierarchical clustering of single cells into distinct cellular subtypes (Wu et al., 2014). Hierarchical clustering yielded only two distinct subsets, which, based on the size distribution of the captured cells, could be identified as LPM (Cluster 1) and SPM (Cluster 2; Fig. 4B). The transcription factor expression profiles of either LPM or SPM showed no further differences between LT-HSC-derived MΦ (RFP^+^) and native MΦ (RFP^−^) (Fig. 4C,D). Similarly, *in vivo* phagocytosis assays, assessing the uptake of *E. coli* by macrophages in the PerC, revealed that both LPM and SPM from either LT-HSCs (RFP^+^) or native origin (RFP^−^) phagocytose bacterial particles at comparable rates (Fig. 4E-G). Collectively, these findings demonstrate that transplanted fetal LT-HSC-regenerated peritoneal MΦ (RFP^+^) are functionally equivalent to their RFP^−^ host counterpart.

**Figure 4.**
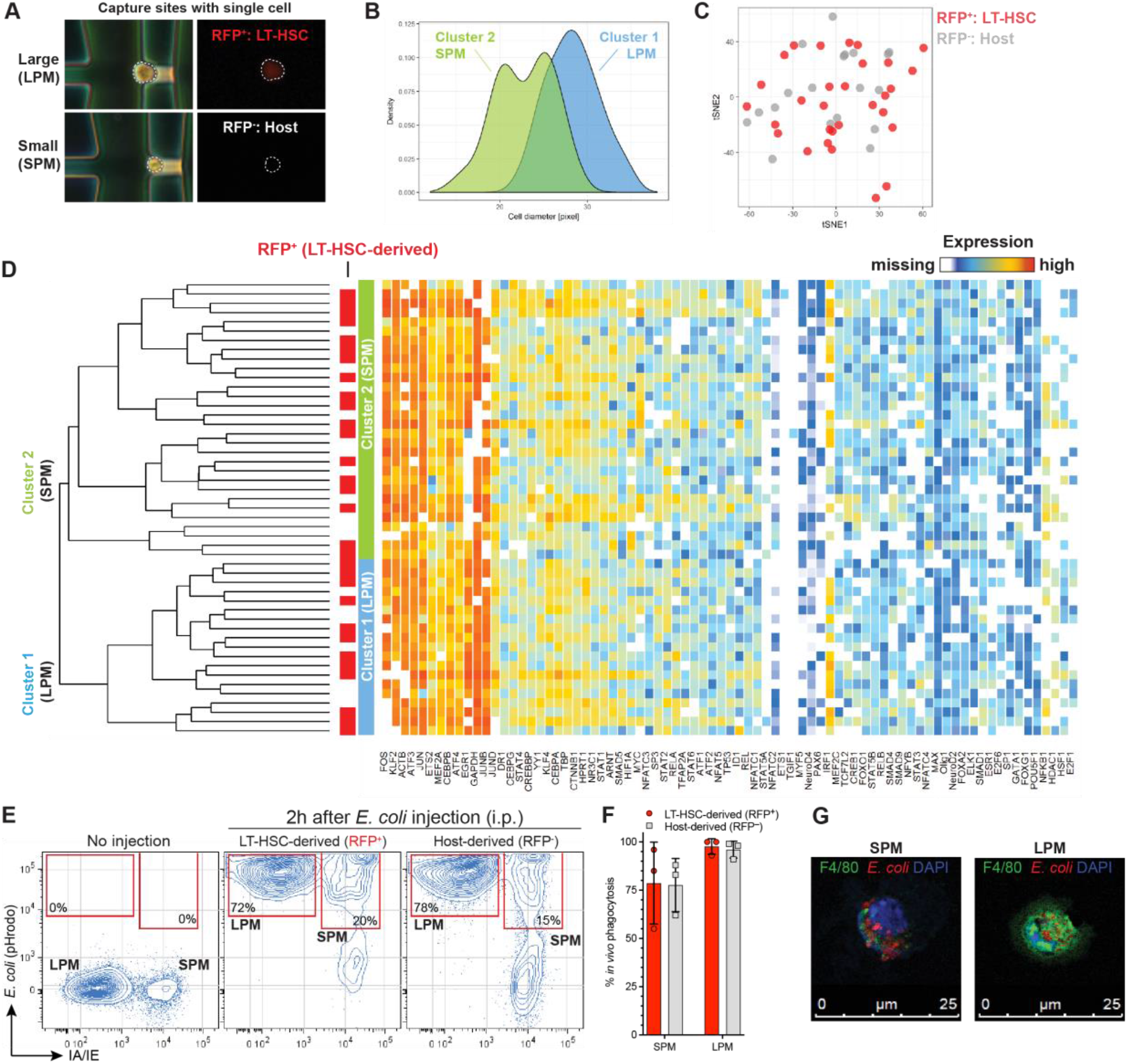
Tissue-resident peritoneal macrophages derived from LT-HSC transplants are functionally comparable to their host-derived counterpart. **A.** Microscopy image of the peritoneal cells after loading them into the Fluidigm C1 fluidics chip. Bright field and fluorescence microscopy images obtained after cell loading reveals the size (small vs. large) and source (RFP^+^ LT-HSC or RFP^−^ host) of the SPM and LPM. **B.** Histograms depicting the size distribution of the two main cell clusters identified by hierarchical clustering of transcription factor expression profiles, shown in D. **C.** t-SNE visualization of the similarity of single isolated peritoneal macrophages derived from transplanted LT-HSC (RFP^+^) or host (RFP^−^), based on transcription factor expression profiles. The lack of defined clusters in t-SNE map indicate that LT-HSC-derived macrophages are similar to their host-derived counterpart. **D.** Hierarchical clustering of transcription factor expression profiles, determined by Fluidigm Biomark single-cell multiplexed qPCR, of single-sorted peritoneal macrophages derived from transplanted LT-HSC (indicated with red bars) or host cells. Analysis yielded two main cell clusters that were identified as LPM (Cluster 1) and SPM (Cluster 2), based on their size distributions shown in B. **E.** Analysis of *in vivo* phagocytosis of pHrodo-labeled *E. coli* particles by LPM (I-A/I-E) and SPM (I-A/I-E^+^) derived from transplanted LT-HSC (RFP^+^) or host cells (RFP), 2 hr after i.p. injection. **F.** Quantification of *in vivo* phagocytosis of *E. coli* particles by LPM and SPM derived from transplanted LT-HSC (RFP^+^, red) or host cells (RFP, gray), as percentage of cells that contained phagocytosed *E. coli* among the specific cell type. **G.** Example of morphology, F4/80 expression and *E. coli* uptake by LPM and SPM. Data shown (E,F) are representative of 2 independent experiments and are mean + SD of 5 total mice (2 control and 3 chimeric animals).

## Discussion

During embryonic development, the first LT-HSC emerges at E10.5, most likely from hemogenic endothelium in the yolk sac and aorta-gonad-mesonephros (AGM) (Hoeffel and Ginhoux, 2015; Kumaravelu et al., 2002; Medvinsky and Dzierzak, 1996; Muller et al., 1994). The current paradigm postulates that these LT-HSCs are capable of continually regenerating all components of the immune system, including the erythroid, lymphoid, and myeloid lineages. However, recent studies have challenged this notion by demonstrating that TRMΦ in the brain (i.e. microglia) and other tissues originate from early progenitors (~E8) that emerge in the yolk sac prior to, and independent of, the development of LT-HSCs (Ginhoux et al., 2010; Gomez Perdiguero et al., 2015a). Though it is becoming increasingly accepted that TRMΦ in the central nervous system (CNS) originate from early yolk sac-derived progenitors that arise before LT-HSCs are developed (Goldmann et al., 2016; Gomez Perdiguero et al., 2015a; Huang et al., 2018; Schulz et al., 2012; Van Hove et al., 2019), there have been conflicting reports as to the origins of TRMΦ populations outside of the CNS (Gomez Perdiguero et al., 2015a; Hoeffel et al., 2015; Sheng et al., 2015a; Yona et al., 2013). In fact, Sheng and colleagues (2015b) proposed that nearly all TRMΦ (with the exception of microglia and a fraction of Langerhans cells) are derived solely from HSCs in the fetal liver, AGM, and/or BM. Here, we demonstrate that both the peritoneum and brain contain CD11b^+^ myeloid immune cells as early as E10 preceding development of the first LT-HSC and *Runx1*^+^ immune progenitor cells labeled at E8 contribute to TRMΦ populations in both the peritoneum and the brain. This suggests that, at a minimum, these populations of TRMΦ in the adult are derived from yolk sac progenitors that emerge from E7.5 until at least E9.5, because they were labeled during a limited time window (approximately 24 h) at E8 (Cline and Moore, 1972; Moore and Metcalf, 1970; Palis et al., 1999; Palis et al., 2001; Samokhvalov et al., 2007). Thus, our lineage tracing experiments demonstrate that tissue-resident LPM and SPM, like CNS-resident microglia, are initially developed from a yolk sac-derived myeloid progenitor that emerges early during embryogenesis and is separate from LT-HSCs, and that these cells persist throughout life.

To directly test the potential of *bona fide* LT-HSCs, from both fetal and adult sources, to regenerate TRMΦ in the peritoneum and brain, we employed adoptive LT-HSC transplantation of solely highly purified LT-HSCs from fetal liver (E15) and adult BM (24 wks). We show that, although the first wave of TRMΦ in the peritoneum and brain originate from the early E8 progenitors, LT-HSC transplants can regenerate peritoneal MΦ but not tissue-resident microglia in the brain. To determine whether LT-HSC-derived MΦ can be distinguished from their host counterparts with regard to their gene expression profile, we performed a targeted single-cell transcriptomic assay (Fluidgim, multiplexed qPCR) on 77 transcription factors. We were not able to detect any significant differences between the transplanted- (RFP^+^) and host- (RFP^-^) derived LPM and SPM, suggesting that purification and transplantation of LT-HSCs does not impact the phenotype and function of the peritoneal MΦ populations that they give rise to. Taken together with the results from our *Runx1* lineage tracing experiments, we provide definitive evidence of *dual ontogeny* for peritoneal MΦ, but not brain microglia. Of note, for the first time, we show that LT-HSCs purified from fetal liver are also unable to generate microglia in the brain. These findings suggest that the microglia population within the CNS does not require LT-HSCs for TRMΦ regeneration, while peripheral tissues, such as the PerC, rely on LT-HSCs to supplement the E8 yolk sac-derived MΦ.

We propose that this difference in sourcing of MΦ progenitors reflect an evolutionary difference in the barrier function of the TRMΦ they give rise to. TRMΦ are an integral part of the body’s first line of defense. However, depending on the tissue, this defense has varying requirements. The PerC is an area of the body that is prone to injury and infection, especially in rodents (Broche and Tellado, 2001; Heemken et al., 1997). Though most TRMΦ regenerate by local self-renewal (Ajami et al., 2007; Ginhoux et al., 2010; Hashimoto et al., 2013; Huang et al., 2018; Jenkins et al., 2011; Yona et al., 2013), sterile injury or inflammation-induced MΦ recruitment (and/or cell death) can rapidly deplete some TRMΦ populations, altering their turnover kinetics (Dannenberg, 2003; Lai et al., 2018; Tay et al., 2017). For example, intestinal MΦ are one of the largest MΦ pools, exhibiting rapid turnover kinetics and reliance on LT-HSC-derived circulating precursors (i.e., monocytes) to maintain normal population densities during homeostasis and inflammation (Bain et al., 2014; Platt et al., 2010). Additionally, cardiac MΦ exhibit a similar trend, where circulating precursors supplement the TRMΦ population with age, even in the absence of inflammation (Epelman et al., 2014; Molawi et al., 2014). Indeed, recent studies have revealed that cavity MΦ, notably LPM and pericardial MΦ, are recruited to injured visceral organs to help mediate clearance of dead/dying cells, promote neovascularization, and prevent fibrosis (Deniset et al., 2019; Gundra et al., 2017; Wang and Kubes, 2016). Therefore, MΦ in these tissues need to be able to regenerate quickly, in order to maintain a sufficient level of protection from infection and/or injury, making them dependent on monocyte-derived MΦ (MDMΦ) in addition to local self-renewal.

In contrast, the CNS has developed in a way that minimizes the risk of inflammation, as bystander damage from inflammatory mediators could have detrimental effects on neurons, which have limited regenerative capacity. The role of microglia is to maintain and restore local homeostasis, only becoming activated if a pathogen does enter the CNS tissue, or, more often, to clear debris from dying CNS-resident cells or prune aberrant synapses (Buttgereit et al., 2016; Li and Barres, 2018; Shemer et al., 2015). They therefore self-renew to replenish, not only because they do not rely on a circulating pool of progenitors, but also because recruiting these cells from circulation could increase the risk of “unwanted” cells or pathogens entering as well. Only when there is a great need to increase MΦ numbers, due to an infection or inflammatory insult in the CNS (e.g., experimental autoimmune encephalitis), will the microglia pool be supplemented with MDMΦ (Ajami et al., 2007; Ajami et al., 2011; Huang et al., 2018). However, infiltrating MDMΦ are unable to differentiate into *bona fide* microglia and remain distinct from the resident microglia population, further supporting the notion that ontogeny, in addition to local microenvironment, influence MΦ heterogeneity (Cronk et al., 2018; Gosselin et al., 2014; Lavin et al., 2014; Shemer et al., 2018; Van Hove et al., 2019).

There is a growing interest in understanding molecular mechanisms that dictate TRMΦ heterogeneity and whether ontogeny influences phenotypic and functional plasticity of these cells. There is a mounting body of evidence that suggests TRMΦ can have divergent responses when compared to infiltrating MDMΦ during inflammatory responses and disease pathogenesis (Ajami et al., 2011; Chen et al., 2019; Lai et al., 2018). In multiple visceral organs, TRMΦ populations that arise independent of LT-HSCs are supplemented or replaced by MDMΦ during inflammation, aging, and/or ablative therapy (Bain et al., 2014; Bain et al., 2016; Bain et al., 2020; Cain et al., 2013; Epelman et al., 2014; Liu et al., 2019; van de Laar et al., 2016). Recruitment of LPM to sites of tissue damage, sterile injury, and/or inflammatory milieus (Deniset et al., 2019; Gundra et al., 2017; Wang and Kubes, 2016) places peritoneal MΦ into a category of TRMΦ that exhibit a dual ontogeny arising from both early yolk sac progenitors and from LT-HSCs. It remains to be determined if these differences in ontogeny influence the functional capacity of MΦ in the peritoneum as evident in MΦ populations (resident vs. infiltrating) in the CNS and other tissues.

In summary, we demonstrate that the requirement for LT-HSCs to regenerate TRMΦ is tissue-specific. Though both peritoneal MΦ and microglia arise from early yolk sac-derived precursors, MΦ in the PerC, but not microglia in the brain, require LT-HSCs to maintain/regenerate their population. We propose that divergence in sourcing of MΦ progenitors, in part, reflects an evolutionary difference in the barrier function of the TRMΦ population they give rise to. These findings add a new layer to the complex developmental landscape of the myeloid lineage, where peritoneal MΦ populations exhibit a definitive dual ontogeny. Though we demonstrate that LT-HSC transplant-derived LPM and SPM are phenotypically and functionally similar to their naturally occurring, host-derived counterparts, whether there is divergence in phenotype and function between those derived from yolk sac progenitors versus fetal and adult LT-HSCs is an outstanding question, warranting further investigation.

## Author contributions

D.E., A.K., H.K., and E.G. designed and performed experiments, analyzed data, prepared and revised the manuscript. J.W., J.S., K.Y., and M.P. performed experiments, reviewed and approved manuscript. E.G. and L.H. provided funding, project administration/supervision, and reagents. All authors discussed the results and contributed to the final manuscript.

## Acknowledgments

We thank Irving L. Weissman (Stanford) for providing pCx-eGFP and TM7-RFP mice, Robert Durruthy-Durruthy (Fluidigm) for his technical direction with the single-cell multiplexed qPCR experiments, and both the Emory Pediatric/Winship Flow Cytometry Core (access supported in part by Children’s Healthcare of Atlanta) and Stanford Shared FACS Facility for their support with flow cytometry experiments. This work was supported by NIH/NIAID R01AI123126 (E.G., Emory), funds from the Lowance Center for Human Immunology (E.G., Emory), and the Herzenberg Laboratory (L.H., Stanford). D.E. was additionally supported by the Laney Graduate School Fellowship (Emory). The funders had no role in study design, data collection and analysis, decision to publish, or preparation of the manuscript.

