## Supplemental Materials for "Hematopoietic stem cell requirement for macrophage regeneration is tissue-specific"

**Table S1. FACS staining reagents used in this study.** Marker, format, isotype, clone, source, and catalog number for all monoclonal antibodies (mAbs) used in fluorescent cytometry experiments for this study. Combinations and staining procedures are listed in experimental methods and FMO controls were used to distinguish auto-fluorescent cells from cells expressing low levels of a particular surface marker as described (see methods).

| Marker | Format | Isotype | Clone | Source | Catalog No. |
|---|---|---|---|---|---|
| B220 | BV786 | Rat IgG2a, κ | RA3-6B2 | BD Biosciences | 563894 |
| B220 | PE-Cy7 | Rat IgG2a, κ | RA3-6B2 | BD Biosciences | 552772 |
| B220 | APC | Rat IgG2a, κ | RA3-6B2 | BD Biosciences | 553092 |
| B220 | AF700 | Rat IgG2a, κ | RA3-6B2 | BD Biosciences | 553092 |
| CD11b | APC-Cy7 | Rat IgG2b, κ | M1/70 | BD Biosciences | 557657 |
| CD11b | BV711 | Rat IgG2b, κ | M1/70 | BD Biosciences | 563168 |
| CD11b | Pacific Blue | Rat IgG2b, κ | M1/70 | Invitrogen | RM2828 |
| CD16/32 | Purified | Rat IgG2b, κ | 2.4G2 | BD Biosciences | 553142 |
| CD19 | BV786 | Rat IgG2a, κ | 1D3 | BD Biosciences | 563333 |
| CD19 | PE-Cy5.5 | Rat IgG2a, κ | 1D3 | Invitrogen | 35-0193-80 |
| CD117 | BV421 | Rat IgG2b, κ | ACK2 | BioLegend | 135124 |
| CD127 | BV605 | Rat IgG2a, κ | A7R34 | BioLegend | 135041 |
| CD135 | Biotin | Rat IgG2a, κ | A2F10 | BioLegend | 135308 |
| CD150 | PE-Cy7 | Rat IgG2a, λ | TC15-12F12.2 | BioLegend | 115914 |
| CD23 | PE | Rat IgG2a, κ | B3B4 | BD Biosciences | 553139 |
| CD23 | Biotin | Rat IgG2a, κ | B3B4 | BD Biosciences | 553137 |
| CD3ε | APC | Armenian Hamster IgG | 145-2C11 | BioLegend | 100312 |
| CD34 | AF700 | Rat IgG2a, κ | RAM-34 | BD Biosciences | 560518 |
| CD38 | PE-Cy5 | Rat IgG2a, κ | 90 | Invitrogen | 15-0381-81 |
| CD41 | PE-Cy5.5 | Rat IgG1, κ | MWReg30 | In-house custom | — |
| CD41 | BV510 | Rat IgG1, κ | MWReg30 | BioLegend | 133923 |
| CD45 | BV570 | Rat IgG2b, κ | 30-F11 | BioLegend | 103136 |
| CD48 | APC-Cy7 | Armenian Hamster IgG | HM48-1 | BioLegend | 103423 |
| CD5 | PE-Cy5 | Rat IgG2a, κ | 53-7.3 | BioLegend | 100610 |
| CX3CR1 | PE | Mouse IgG2a, κ | SA011F11 | BioLegend | 149006 |
| F4/80 | PE-Cy7 | Rat IgG2a, κ | BM8 | BioLegend | 123114 |
| Gr-1 | APC | Rat IgG2b, κ | RB6-8C5 | BioLegend | 108412 |
| Gr-1 | AF700 | Rat IgG2b, κ | RB6-8C5 | BD Biosciences | 557979 |
| Gr-1 | APC-Cy7 | Rat IgG2b, κ | RB6-8C5 | BD Biosciences | 557661 |
| I-A/I-E | BV650 | Rat IgG2b, κ | M5/114.15.2 | BD Biosciences | 563415 |
| IgK | Biotin | Rat IgG1, κ | 187.1 | BD Biosciences | 559750 |
| IgK | APC-Cy7 | Rat IgG1, κ | 187.1 | BD Biosciences | 561353 |
| IgM | AF700 | | 331 | In-house custom | — |

| | | | | | |
|---|---|---|---|---|---|
| Ly-6C | APC | Rat IgM, κ | AL-21 | BD Biosciences | 560595 |
| Sca-1 | PE | Rat IgG2a, κ | D7 | BioLegend | 108108 |
| Sca-1 | PE-Cy5.5 | Rat IgG2a, κ | D7 | Invitrogen | MSCA18 |
| Streptavidin | BV711 | — | — | BioLegend | 405241 |
| Streptavidin | Qdot605 | — | — | Invitrogen | Q10101MP |
| TER-119 | PE | Rat IgG2b, κ | TER119 | BD Biosciences | 553673 |
| TER-119 | APC | Rat IgG2b, κ | TER119 | BioLegend | 116212 |
| Viability dye | Zombie Aqua | — | — | BioLegend | 423102 |
| Vδ6.3/2 | PE | Armenian Hamster IgG2, κ | 8F4H7B7 | BD Biosciences | 555321 |
| VH11 | Pacific Blue | | VH11Id.6e9 | In-house custom | — |

**Table S2. Fluidigm plate design for single cell HT-qPCR assay.** Target, RefSeq accession number, and gene symbol for the 77 targets used in Fluidigm single-cell HT-qPCR assay (in alphabetical order).

| Target | Design RefSeq | Gene Symbol |
|---|---|---|
| Actb | NM_007393.3 | Actb |
| Arnt | NM_009709.4 | Arnt |
| Atf1 | NM_007497.3 | Atf1 |
| Atf2 | NM_009715.2 | Atf2 |
| Atf3 | NM_007498.3 | Atf3 |
| Atf4 | NM_009716.2 | Atf4 |
| Cebpa | NM_007678.3 | Cebpa |
| Cebpb | NM_009883.3 | Cebpb |
| Cebpg | NM_009884.3 | Cebpg |
| Creb1 | NM_001037726.1 | Creb1 |
| Crebbp | XM_006521751.3 | Crebbp |
| Ctnnb1 | NM_007614.3 | Ctnnb1 |
| Dr1 | NM_026106.3 | DR1 |
| E2f1 | NM_007891.4 | E2f1 |
| E2f6 | NM_033270.1 | E2F6 |
| Egr1 | NM_007913.5 | Egr1 |
| Elk1 | NM_007922.4 | Elk1 |
| Esr1 | NM_007956.4 | Esr1 |
| Ets1 | NM_001038642.1 | Ets1 |
| Ets2 | NM_011809.3 | Ets2 |
| Fos | NM_010234.2 | Fos |
| Foxa2 | NM_010446.2 | Foxa2 |
| Foxg1 | NM_001160112.1 | Foxg1 |
| Foxo1 | NM_019739.3 | Foxo1 |
| Gapdh | NM_008084.2 | Gapdh |
| Gata1 | NM_008089.1 | Gata1 |

| | | |
|---|---|---|
| Hdac1 | NM_008228.1 | Hdac1 |
| Hif1a | NM_010431.2 | Hif1a |
| Hprt | NM_013556.1 | HPRT1 |
| Hsf1 | NM_008296.2 | Hsf1 |
| Id1 | NM_010495.2 | ID1 |
| Irf1 | NM_008390.2 | Irf1 |
| Jun | NM_010591.2 | Jun |
| Junb | NM_008416.2 | Junb |
| Jund | NM_010592.4 | Jund |
| Klf2 | NM_008452.2 | Klf2 |
| Klf4 | NM_010637.3 | Klf4 |
| Max | NM_008558.1 | Max |
| Mef2a | NM_001033713.1 | Mef2a |
| Mef2c | NM_001170537.1 | Mef2c |
| Myc | NM_010849.4 | Myc |
| Myf5 | NM_008656.5 | Myf5 |
| Neurod2 | NM_010895.2 | Neurod2 |
| Neurod4 | NM_007501.4 | Neurod4 |
| Nfat5 | NM_018823.2 | Nfat5 |
| Nfatc1 | NM_016791.4 | Nfatc1 |
| Nfatc3 | NM_010901.2 | Nfatc3 |
| Nfatc4 | NM_023699.3 | Nfatc4 |
| Nfkb1 | NM_008689.2 | Nfkb1 |
| Nfyb | NM_010914.1 | Nfyb |
| Nr3c1 | NM_008173.3 | Nr3c1 |
| Olig1 | NM_016968.3 | Olig1 |
| Pax6 | NM_013627.4 | Pax6 |
| Pou5f1 | NM_013633.2 | Pou5f1 |
| Rel | NM_009044.2 | Rel |
| Rela | NM_009045.4 | Rela |
| Relb | NM_009046.2 | Relb |

| | | |
|---|---|---|
| Smad1 | NM_008539.3 | Smad1 |
| Smad4 | XM_001001668.1 | Smad4 |
| Smad5 | NM_008541.3 | Smad5 |
| Smad9 | NM_019483.4 | Smad9 |
| Sp1 | NM_013672.2 | Sp1 |
| Sp3 | NM_001098425.1 | Sp3 |
| Stat1 | NM_009283.3 | Stat1 |
| Stat2 | NM_019963.1 | Stat2 |
| Stat3 | NM_213659.2 | Stat3 |
| Stat4 | NM_011487.4 | Stat4 |
| Stat5a | NM_011488.3 | Stat5a |
| Stat5b | NM_011489.2 | Stat5b |
| Stat6 | NM_009284.2 | Stat6 |
| Tbp | NM_013684.3 | Tbp |
| Tcf7l2 | NM_001142918.1 | Tcf7l2 |
| Tcf7l2 | NM_009333.3 | Tcf7l2 |
| Tfap2a | NM_011547.3 | Tfap2a |
| Tgif1 | NM_009372.2 | Tgif1 |
| Trp53 | NM_011640.2 | Tp53 |
| Yy1 | NM_009537.2 | Yy1 |

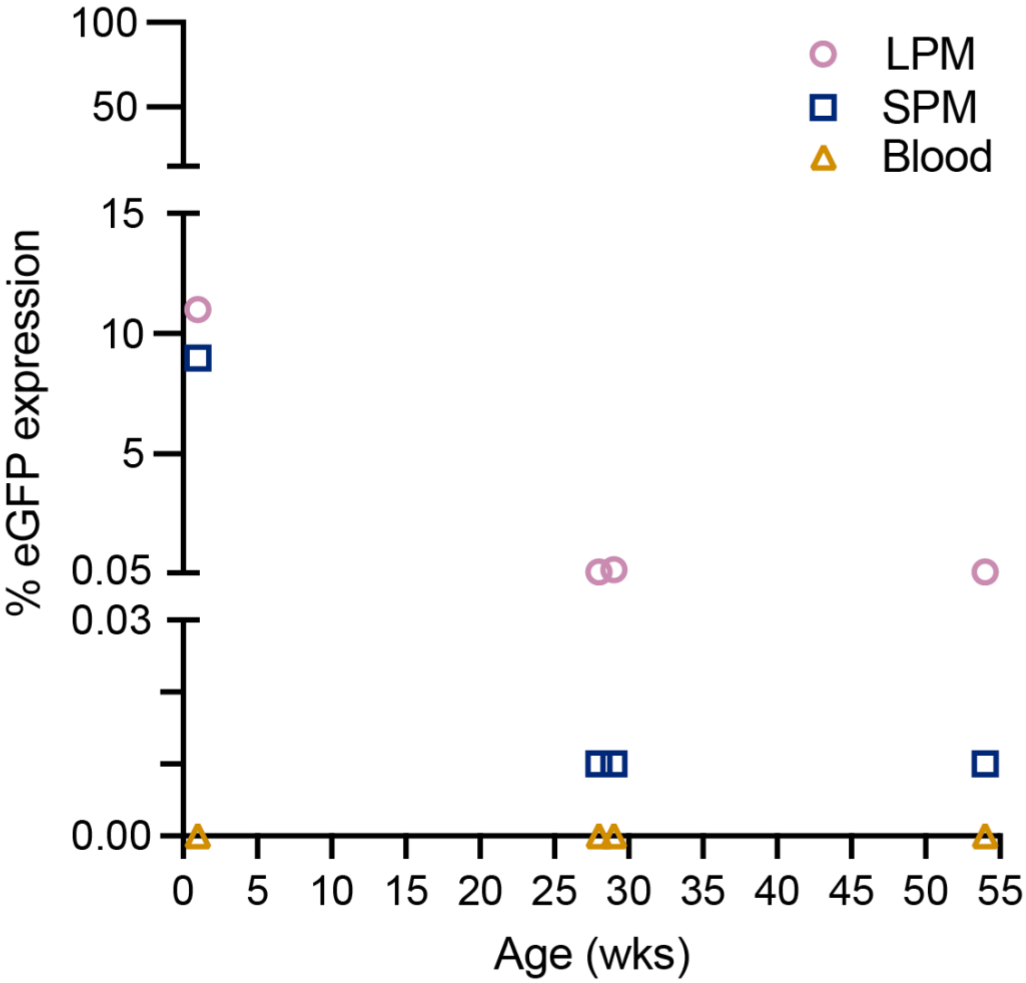


**Figure S1. Frequency of eGFP labeled cells in *Runx1* lineage tracing mice at age of sacrifice.** Each individual *Runx1*$^{cre/eGFP}$ mouse shown in Fig. 2 is plotted at age of sacrifice to show higher proportions of eGFP cells in the peritoneal cavity of young mice (1 wk old) that declines with age yet persists suggesting some of the yolk-sac-derived population is likely maintained by self-renewal of eGFP-expressing LPM and SPM.

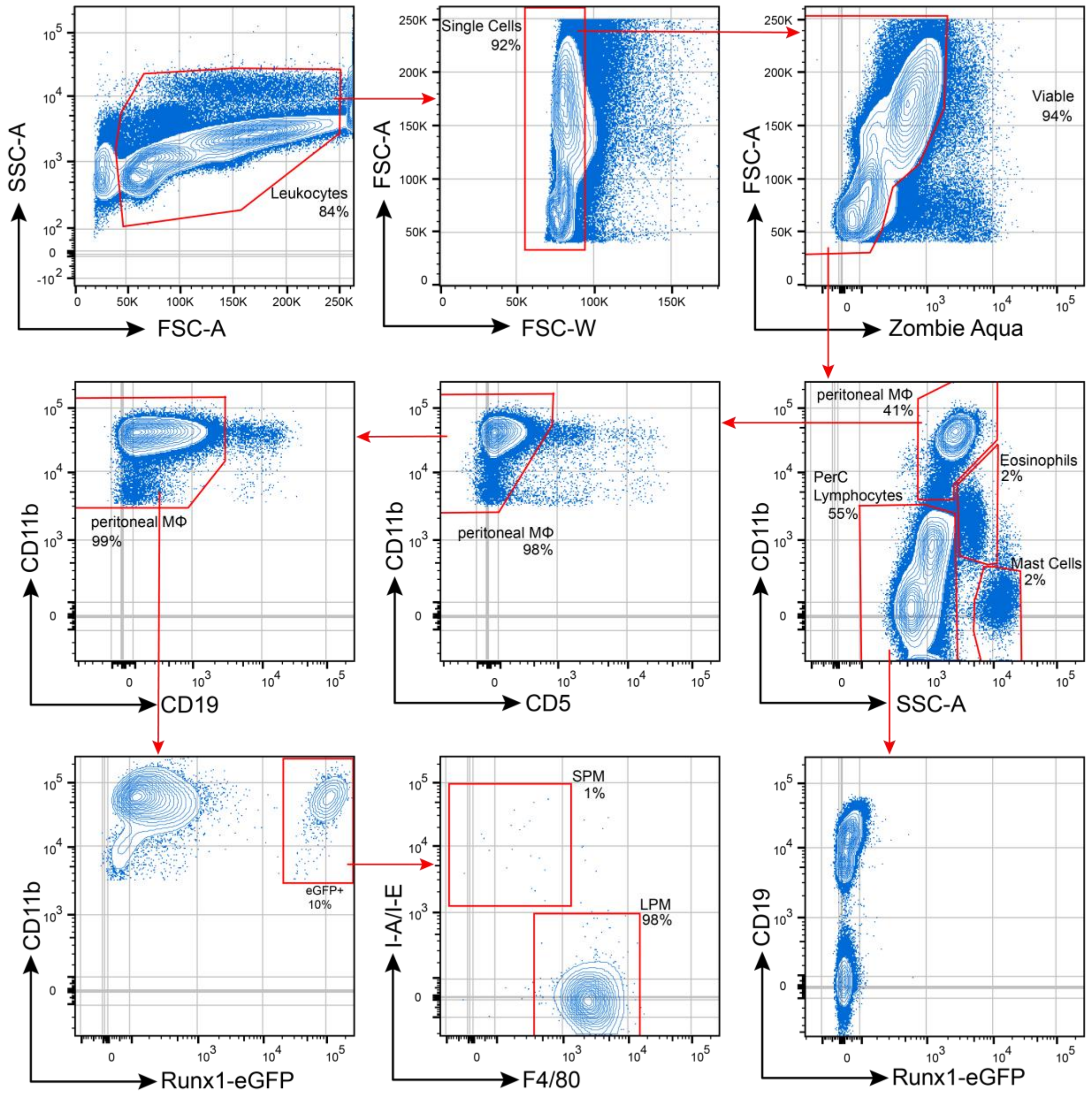


**Figure S2. Representative gating strategy–Peritoneal Cavity (PerC).** Viable, single cells were gated first using CD11b and SSC-A to identify major immune lineages in the PerC as previously established (Ghosn et al. 2010). Gated peritoneal macrophages (MΦ) were "cleaned: for T/B cells doublets (removing CD5 and CD19, respectively) then fluorescent reporters interrogated in lineage tracing (*Runx1*$^{cre/eGFP}$) or transplanted mice (RFP & GFP) as shown in Figs. 2 & 3.  SPM and LPM were distinguished using I-A/I-E (MHC II) and F4/80 expression as previously established (Ghosn et al. 2010). Gated PerC lymphocytes were interrogated for eGFP expression in *Runx1*$^{cre/eGFP}$ mice to confirm that LT-HSCs were not labeled.

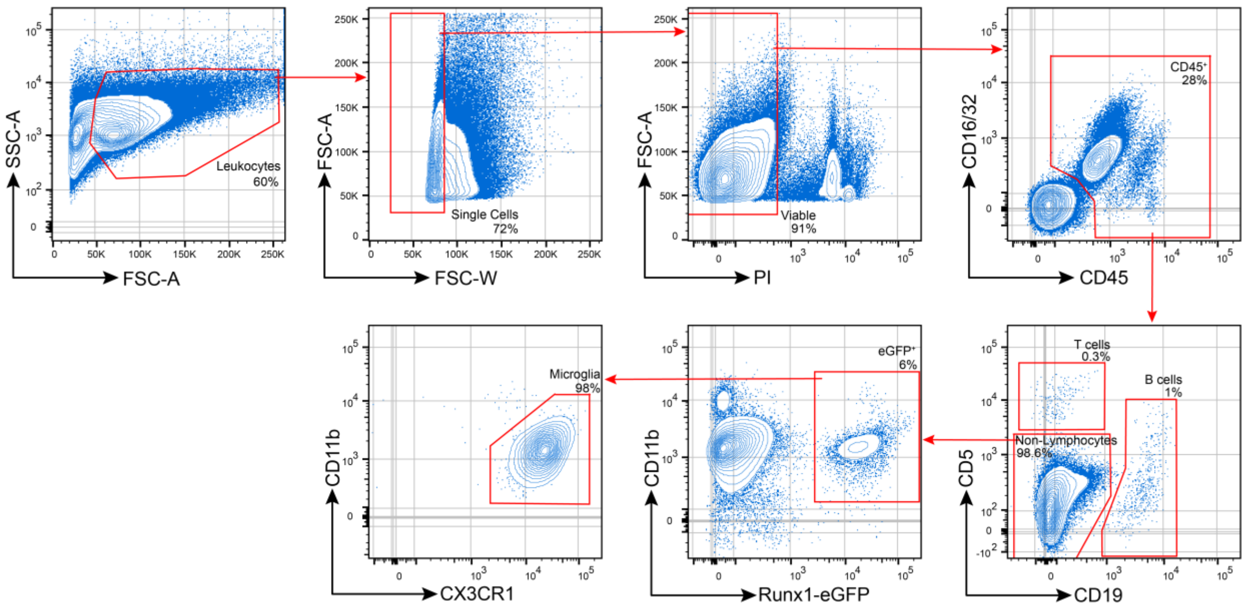


**Figure S3. Representative gating strategy–Brain.** Viable, single cells were gated for CD45+ cells to analyze immune populations in the brain. CD5+ T cells and CD19+ B cells were gated and remaining CD11b+ cells analyzed for *Runx1*$^{cre/eGFP}$ expression and RFP/GFP expression in transplanted mice in gates displayed in Figs. 2 & 3.

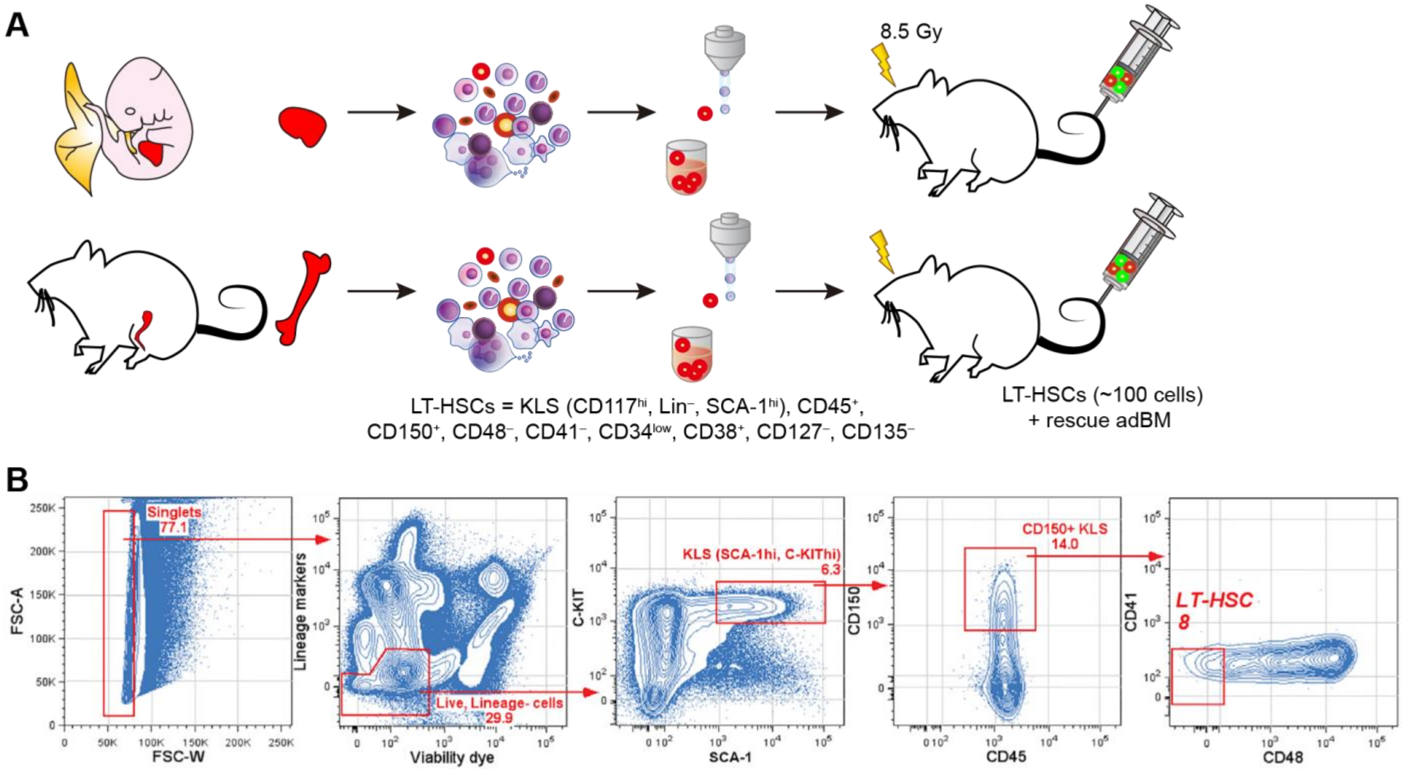


**Figure S4. LT-HSC adoptive transplantation strategy. A.** Fetal liver (E15) and adult BM (adBM) are isolated from TM7-RFP mice and single LT-HSCs sorted as described (see methods) for adoptive transplantation into lethally-irradiated recipient mice. Mice also received GFP$^{+}$ adBM rescue cells as described at the time of LT-HSC transplantation (see methods). **B.** Representative sorting strategy for RFP$^{+}$ LT-HSCs from fetal (shown here) and adBM. Adapted from Ghosn et al. (2016).

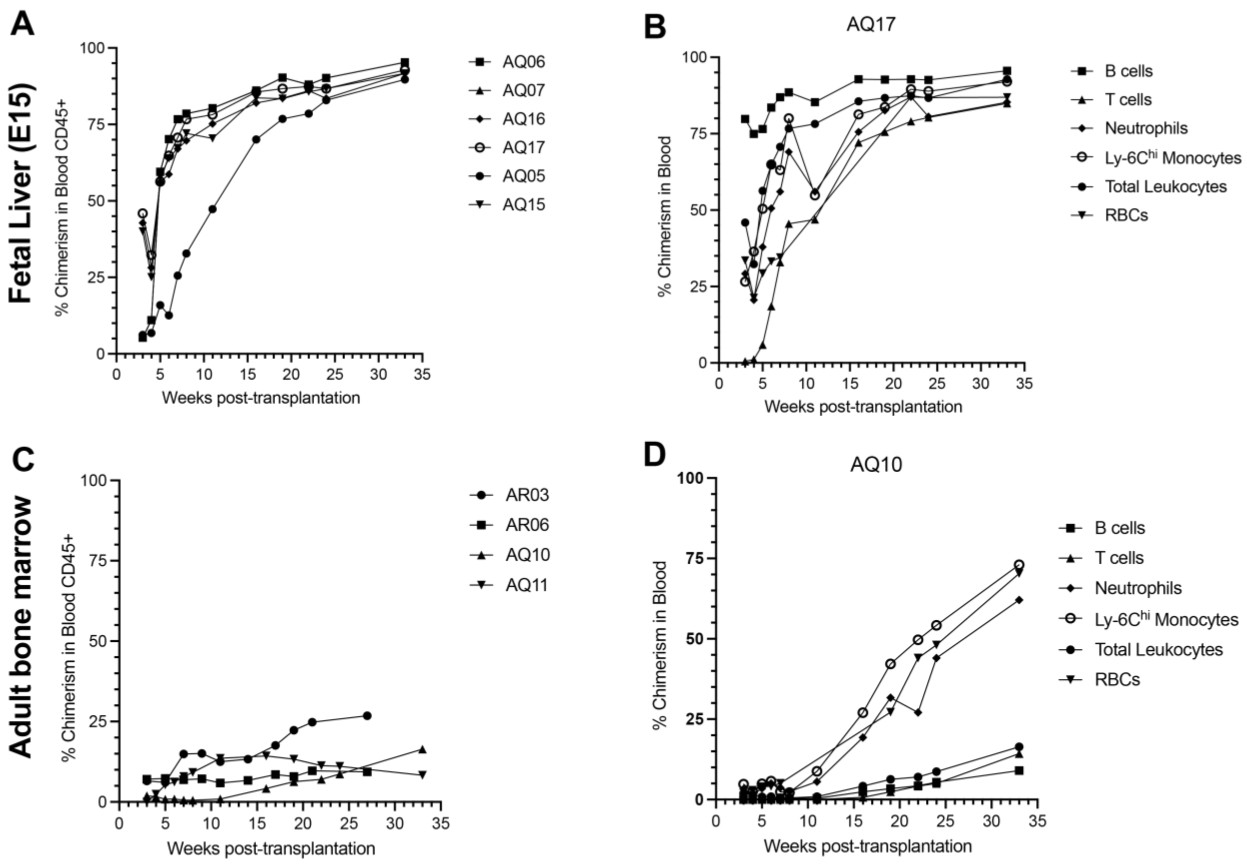


**Figure S5. Representative blood RFP chimerism kinetics of transplant recipient mice. A.** Chimerism kinetics of total CD45+ blood leukocytes for 6 representative mice that received E15 fetal liver (FL) RFP+ LT-HSC transplants. **B.** Chimerism kinetics of major immune lineages from one representative mouse that received E15 FL RFP+ LT-HSC transplant. **C.** Chimerism kinetics of total CD45+ blood leukocytes for 4 mice that received adult bone marrow (adBM) RFP+ LT-HSC transplant. **D.** Chimerism kinetics of major immune lineages from one representative mouse that received adBM RFP+ LT-HSC transplant.
